# Temporal structure of item presentation modulates brain oscillations in verbal working memory

**DOI:** 10.1101/2025.05.04.652111

**Authors:** Alexandra I. Kosachenko, Danil I. Syttykov, Dmitrii A. Tarasov, Alexander I. Kotyusov, Dauren Kasanov, Sergey Malykh, Boris Kotchoubey, Yuri G. Pavlov

## Abstract

Previous studies of verbal working memory (WM) have reported inconsistent changes in alpha power during retention, with both increases and decreases observed. We asked whether these discrepancies arise from how stimuli are presented. Thirty adults memorized seven digits presented in four modes: Simultaneous (all digits for 2800 ms) or sequential presentations at Fast (400 ms per digit), Slow (1000 ms per digit), and Fast+delay (400 ms per digit plus a 600 ms free time in between). We analyzed EEG during encoding and a 6 s retention period in theta (4-7 Hz), alpha (8-13 Hz), and beta (18-24 Hz) frequency bands. Encoding produced parametric load-related theta increase and beta decrease, possibly reflecting growing executive control demands and motor program formation, respectively. Alpha power did not scale with load during encoding. Single trial models linked stronger encoding theta and deeper alpha suppression to better recall, whereas retention power did not predict accuracy. During retention, theta and beta were unaffected by presentation mode. Alpha power did not differ significantly when all sequential modes were grouped together compared to the simultaneous mode. However, the Fast+delay mode uniquely showed below-baseline alpha in the first half of the retention. Our findings suggest that alpha dynamics are sensitive to the temporal structure of encoding and retention periods, particularly the presence or absence of free intervals between stimulus presentations. We propose that alpha modulation during WM retention may reflect processes beyond the simple gating of irrelevant sensory information.

## 1 Introduction

Working memory (WM) is the ability to maintain and manipulate information over short durations (Hitch et al., 2024), playing a crucial role in everyday cognition, including higher-order functions such as intelligence and language (Gathercole, 2012; Gignac, 2014). Brain oscillations appear to support the encoding and maintenance of WM. In particular, alpha activity (8-13 Hz) has frequently been implicated in WM functioning, with alpha power varying based on how many items are held in memory during the retention period (Pavlov & Kotchoubey, 2022).

The early discovery that alpha activity during WM retention sometimes increases with load (Jensen et al., 2002) prompted researchers to move away from viewing the alpha rhythm as merely reflecting “cortical idling”. Instead, the gating-by-inhibition hypothesis (Jensen & Mazaheri, 2010) posits that alpha activity serves as a functional inhibitor of irrelevant cortical areas, thus optimizing activity in task-critical regions. From this perspective, alpha enhancement during verbal WM retention corresponds to inhibiting the visual sensory cortex, which is not actively involved in verbal stimulus processing. In contrast, retaining visuospatial information requires ongoing engagement of the visual sensory cortex, and alpha power may drop below baseline (Van Ede, 2018). At the same time, alpha increases have been observed alongside strong sensory responses, as cross modal frequency tagging shows larger visual and auditory steady state responses when alpha is elevated, a pattern more consistent with alpha shaping later information routing than with simple suppression of early sensory gain (Brickwedde et al., 2025). Nevertheless, in his most recent update to the inhibition-by-alpha theory, Jensen (2024) reinforces the idea that alpha power decreases when complex visuospatial information is maintained, yet increases with growing WM load during the retention of letters, consonants, or faces. However, this pattern has been replicated in many but not in all studies. A systematic review examining tasks where encoding and retention are temporally separated (e.g., Sternberg tasks, simple span tasks) found that alpha enhancement during retention occurred in approximately 80% of verbal WM studies and around 60% of those with a visual modality of the stimuli (Pavlov & Kotchoubey, 2022). Thus, while stimulus modality matters, it alone does not explain the variability in previous findings. Moreover, inconsistencies even within the verbal domain raise an important question - which additional factors contribute to the variability of alpha activity during verbal WM retention?

Task features can influence whether alpha power is enhanced or suppressed during the retention period of a WM task. Previous studies from our group have shown both alpha enhancement (Kasanov et al., 2024; Pavlov & Kotchoubey, 2020) and alpha suppression (Kosachenko et al., 2023) during verbal WM retention. One major distinction among these studies was the mode of item presentation: memory items were presented either simultaneously (Kasanov et al., 2024; Pavlov & Kotchoubey, 2020) or sequentially (Kosachenko et al., 2023). These findings suggest that differences in alpha activity may stem from how memory items are presented. Although a systematic review reported no significant differences in the prevalence of alpha enhancement vs. suppression between sequential and simultaneous presentation (Pavlov & Kotchoubey, 2022), only one experiment made a direct comparison. In that experiment, Okuhata et al. (2013)) found decreased alpha power during retention in the simultaneous condition of a Sternberg task and increased alpha in the sequential condition. However, shorter time was available for encoding in the simultaneous condition compared to the sequential condition, which may have further influenced alpha dynamics. Additionally, their presentation of verbal stimuli was unconventional because items were placed in a circle around the fixation cross, potentially disrupting the typical temporal structure of the memoranda. This setup may have emphasized visual rather than verbal strategies. Chen et al. (2022), who used a similar format, also observed strong alpha suppression during retention for both verbal and visual stimuli. They proposed that posterior alpha power increases with WM load when tasks require verbal strategies but decreases when tasks require maintaining visual identity. Collectively, these findings suggest that the decades-long variability reported in alpha activity during WM retention may partly stem from differences in presentation mode.

In sequential presentation mode, commonly employed in experimental psychology but less so in cognitive neuroscience, the duration of memory item presentation has been shown to significantly influence behavioral performance. Longer item presentation, up to about 4 s (Unsworth, 2016), improves memory accuracy. Moreover, the amount of free time between memory items (i.e., the blank interval following the encoding of each sequentially presented item) plays a critical role in preserving the memoranda. Recent studies have reported improved long-term memory for an item held in WM when additional free time is provided after its initial presentation (Hartshorne & Makovski, 2019; Jarjat et al., 2018, 2020; Souza & Oberauer, 2017). One plausible explanation is that, in short presentation conditions, rehearsal is postponed until the retention interval, whereas in long-presentation conditions, rehearsal can occur between items. At least one study has explicitly linked this advantage of additional free time to increased WM consolidation (Cotton & Ricker, 2021). These consolidation differences between presentation modes may affect performance and could also be reflected in neural correlates of WM retention and encoding. However, to date, no EEG studies have directly tested this hypothesis.

Whereas the direction of alpha changes related to WM maintenance remains disputed, there is less controversy concerning theta activity. In sequential presentation studies, load effects can be observed at the level of individual memory items (Kosachenko et al., 2023; Onton et al., 2005) or manifest as a parametric increase in frontal theta power with a greater overall number of WM items during retention interval when items are presented simultaneously (Jensen et al., 2002b; Scheeringa et al., 2009). The prevailing explanation is that theta increases with the growing need for executive control over sensory cortical areas (Ratcliffe et al., 2022). However, there is insufficient evidence to determine whether presentation mode - simultaneous vs. sequential - affects theta, or whether the loading function might differ under shorter vs. longer stimulus-onset asynchrony (SOA) when potentially less time is available to exert executive control.

Beta activity is rarely studied but has likewise been implicated in WM maintenance (Berger et al., 2014; Erickson et al., 2017; Pavlov & Kotchoubey, 2021; Wen et al., 2024) and, more recently, in executive control (Liljefors et al., 2024). In serial recall tasks, it is reasonable to hypothesize that motor planning, reflected by beta suppression over motor cortical areas, may also be involved (Nasrawi et al., 2025). Presentation mode, in turn, may affect the likelihood of encoding incorrect motor programs, or shorter SOAs may constrain motor planning by consuming resources needed for sensory encoding - both factors could contribute to differences in beta activity across conditions.

In the present study, we examined how different presentation modes affect EEG correlates of WM encoding and retention by comparing four conditions in which sets of digits were presented: (i) simultaneously, (ii) sequentially with a short presentation time (400 ms), (iii) sequentially with a long presentation time (1000 ms), and (iv) sequentially with a 600 ms blank interval after each item. This design allowed us to systematically assess how alpha, theta, and beta activity vary as a function of timing, inter-stimulus intervals, and simultaneous vs. sequential presentation. Based on the current theoretical framework, we hypothesized that alpha power during the retention interval would increase relative to baseline across all presentation modes. We also explored, without strong a priori predictions, whether frontal midline theta and lower beta activity exhibit load-dependent changes in the sequential conditions and tested whether these patterns of changes differ from those observed under simultaneous presentation.

## 2 Methods

### 2.1 Participants

Thirty-five native Russian speakers initially participated in the experiment. Five participants were excluded due to technical issues during EEG recording, resulting in a final sample of thirty participants (25 females, mean age ± SD: 19.9 ± 2.7 years). Of the final sample, 28 were recruited from Ural Federal University (27 from the Department of Psychology and 1 from another department) and 2 from the local community. All participants had normal or corrected-to-normal vision and were free from any current neurological or psychiatric disorders. Informed consent was obtained from each participant. The experimental protocol was approved by the Ural Federal University Ethics Committee.

### 2.2 Task

Participants completed four blocks of a WM task, each corresponding to one of four presentation modes (see Figure 1) and consisting of 50 trials. The order of these blocks was randomized across participants. Each trial began with an exclamation mark displayed on the screen for a random duration of 1-2 s, followed by a baseline fixation cross (Thaler et al., 2013) for 2 s. Next, participants were presented with seven unique digits for encoding, after which they fixated the cross and retained the digits in memory for 6 s. In Simultaneous mode, digits were shown simultaneously for 2800 ms. In the other three conditions, digits were presented sequentially: in Fast mode, each digit appeared for 400 ms; in Slow mode, for 1000 ms; and in Fast+delay mode, for 400 ms with a 600 ms interstimulus interval while the fixation cross remained on-screen. In the latter condition, the retention period was reduced to 5.4 s to keep the overall retention period constant across conditions. Immediately after the retention, participants entered the digits in the correct serial order via a numeric keypad. Feedback was provided only during an 8-trial training session that preceded the main task phase.

**Figure 1.**
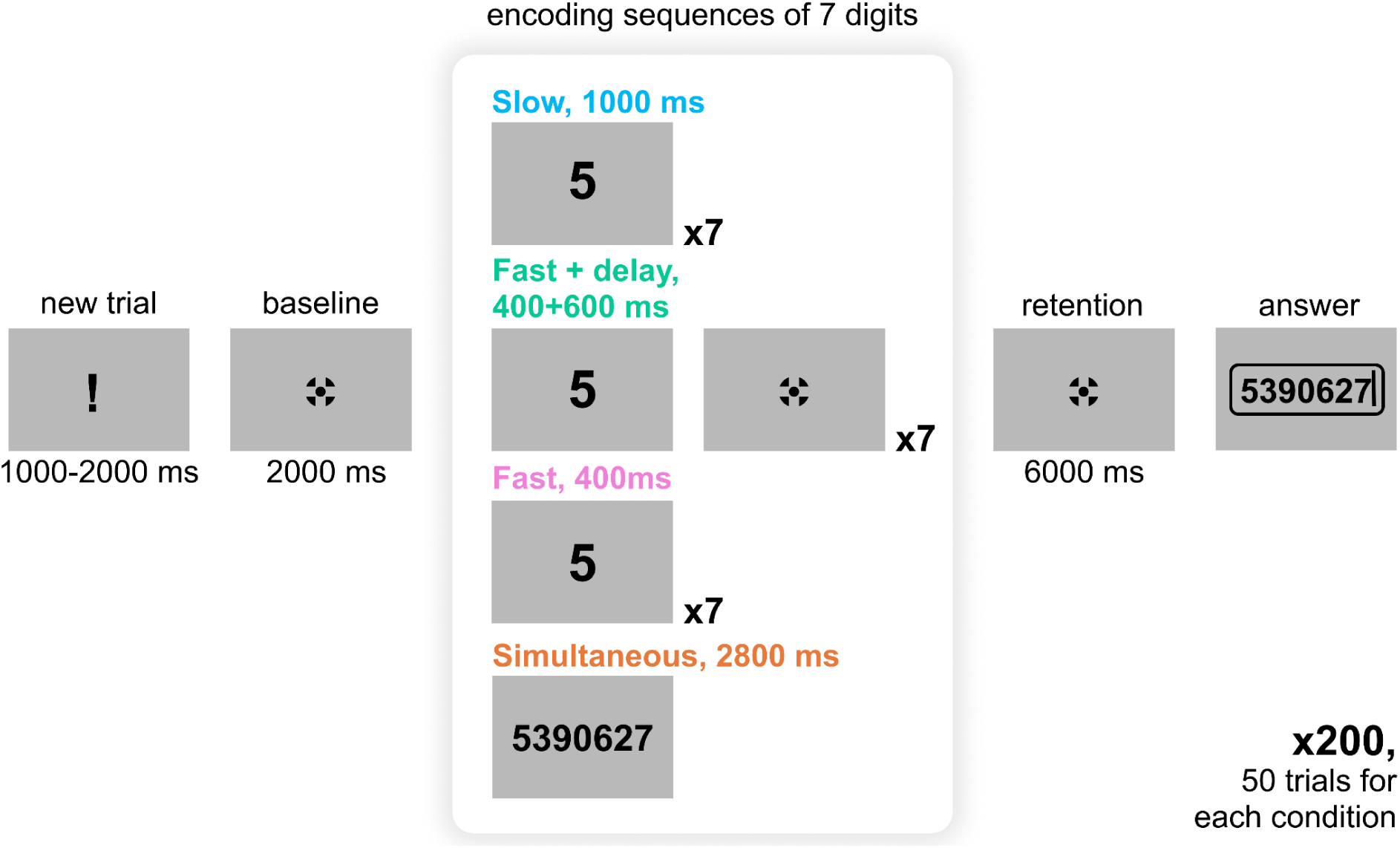
Task design (not to scale). Participants memorized a sequence of 7 digits presented in one of four distinct modes: Simultaneous presentation for 2800 ms, or Sequential presentation in Fast (400 ms per digit), Slow (1000 ms per digit), and Fast+Delay (400 ms per digit with a 600 ms pause) modes. Following the encoding period, participants retained the digits in memory for 6 seconds before recalling the entire set in the correct serial order.

Participants received self-paced breaks every 25 trials and three longer breaks between blocks, with drinks and snacks available on demand. On average, the experiment lasted 94 min. The task was developed using PsychoPy 2023.1.2 (Peirce et al., 2019).

### 2.3 Behavior analysis

As a primary measure of behavioral accuracy, we calculated partial score as the number of digits recalled in the correct serial position.

We also analyzed the time participants took to enter their responses, treating this as a measure of retrieval time. Participants could correct their entries using the backspace key. To ensure consistent measurement of retrieval duration (input time) for both the entire response and each digit, we only included trials in which participants submitted their answers without any corrections (including those due to accidental key presses). This restriction did not apply to reaction time (the interval before typing the first digit).

### 2.4 Electroencephalography

A 64-channel EEG system with active electrodes (ActiCHamp, Brain Products) was used for the recording. The electrodes were placed according to the extended 10-20 system with the FCz channel as the online reference and Fpz as the ground electrode. The level of impedance was maintained below 25 kOm. The sampling rate was 1000 Hz.

EEGLAB (Delorme & Makeig, 2004) was used for data preprocessing. Each recording was filtered by applying 1 Hz high-pass and 45 Hz low-pass filters. After this, data were downsampled to 250 Hz, re-referenced to the average reference, and the Independent Component Analysis was performed using the AMICA algorithm (Palmer et al., 2011). Components clearly related to eye movements were removed. Additionally, components containing channel noise or muscle activity that were mapped onto one electrode and could be clearly distinguished from EEG signals were subtracted from the data. The data were epoched in [−10000 to 8000 ms] intervals in relation to the retention interval onset. These extended epochs were used to minimize edge artifacts during subsequent analyses. Any remaining epochs containing artifacts were manually inspected and discarded.

In Fieldtrip toolbox (Oostenveld et al., 2011), time-frequency analysis was performed on the preprocessed single trial data between 1 and 45 Hz with 1 Hz steps using Morlet wavelets with the number of cycles varying from 3 to 12 in 45 logarithmically spaced steps for each participant and condition, separately. The analysis time window was shifted in steps of 20 ms. Spectral power was baseline-normalized by computing the percent change of the power with respect to the [−1000 to −100] ms time interval, which corresponded to 900 ms of the baseline fixation before the presentation of the stimulus set.

Three frequency bands and regions of interest (ROIs) were identified: frontal midline theta (4-7 Hz, channel Fz), central lower beta (18-24 Hz, channel C3), and posterior alpha (9-13 Hz, channels PO7, PO3, POz, PO4, PO8, O1, Oz, and O2). ROI definitions were guided by prior studies and by inspection of the flattened average time frequency maps across participants and conditions (Bowman et al., 2020). Fz was selected for quantifying frontal midline theta due to its widespread use in previous studies (Kosachenko et al., 2023; Pavlov & Kotchoubey, 2020). C3 was chosen because it exhibited the most pronounced beta suppression on flattened average and as a canonical electrode for motor working memory (Boettcher et al., 2021). The alpha ROI targeted occipito-parietal cortex where robust delay period alpha was observed. We analyzed lower beta only because task related beta effects in this dataset were confined to 18-24 Hz, whereas higher beta showed no consistent modulation. To reduce cross talk between bands, CP electrodes were excluded from both alpha and beta ROIs. An additional analysis using all channels is reported in Supplementary Figure S1.

### 2.5 Statistics

A repeated-measures (rm) ANOVA was performed on behavioral accuracy with two factors: Presentation mode (Simultaneous, Fast, Slow, or Fast + delay presentation modes) and Serial position (digits 1-7). The same approach was used to analyze retrieval time. In the post-hoc t-tests, all p-values were adjusted using the Benjamini-Hochberg (BH) procedure to control the false discovery rate (Benjamini & Hochberg, 1995), and Greenhouse-Geisser corrections were applied to the degrees of freedom where necessary.

For the EEG analyses, the retention interval was divided into two halves, consistent with our previous study (Pavlov & Kotchoubey, 2021) and to align with typically shorter retention intervals in other comparative studies. The first 0.6 seconds of Retention were excluded to eliminate the effect of stimulus change. The spectral power data were averaged across three time windows (periods): Encoding, Retention 1 (0.6-3 s), and Retention 2 (3-6 s). The ANOVA for these data included Period (three levels) and Presentation mode.

Next, rather than using an ANOVA that does not effectively capture linear trends, we explored the linear effect of Serial position (WM load) on the spectral power in the three frequency bands and tested whether the slope of this change differed across presentation modes using linear regression analysis. Relative spectral power was averaged over intervals corresponding to the encoding and maintenance of each digit (400 ms for Fast, 1000 ms for Slow and Fast + delay modes, or equivalent intervals in the Simultaneous mode. Specifically, for Simultaneous mode, the encoding period was segmented into seven 400-ms intervals, mirroring the Fast presentation mode. This analysis included Serial position (seven levels) and Presentation mode (four levels: Simultaneous, Fast, Fast + delay, Slow).

To link behavior and EEG within participants, we tested whether single-trial spectral power predicted the number of correctly recalled digits. We fit generalized mixed-effects models with recall count 0-7 as the outcome and baseline-normalized spectral power and presentation mode as predictors. Power was centered. Presentation mode was contrast coded. Each model included random intercepts and random slopes for spectral power and presentation mode by participant. We ran separate models for each frequency band and task period. As a robustness check, we repeated the encoding analyses after excluding the first 400 ms of encoding.

All statistical analyses were conducted in R (v. 4.3.2).

## 3 Results

### 3.1 Behavior

Presentation mode had a significant effect on recall accuracy (main effect of Presentation mode: F(2.68, 77.77) = 59.92, p < .001, *η_p_²* = .67; see Figure 2). Pairwise comparisons showed significant differences between all conditions (all p < .001, except between Slow and Fast + Delay presentations with p = .025). Participants recalled the most digits when they were presented simultaneously (mean ± SD = 6.27 ± 1.31) and the fewest in the Fast sequential presentation mode (400 ms; 4.79 ± 2.15). When the SOA was kept constant at 1 second but encoding time varied (1000 ms for Slow versus 400 ms for Fast + Delay), recall performance remained significantly different, although the magnitude of the difference was smaller, with averages of 5.74 ± 1.76 digits for Slow and 5.52 ± 1.88 digits for Fast + Delay conditions.

**Figure 2.**
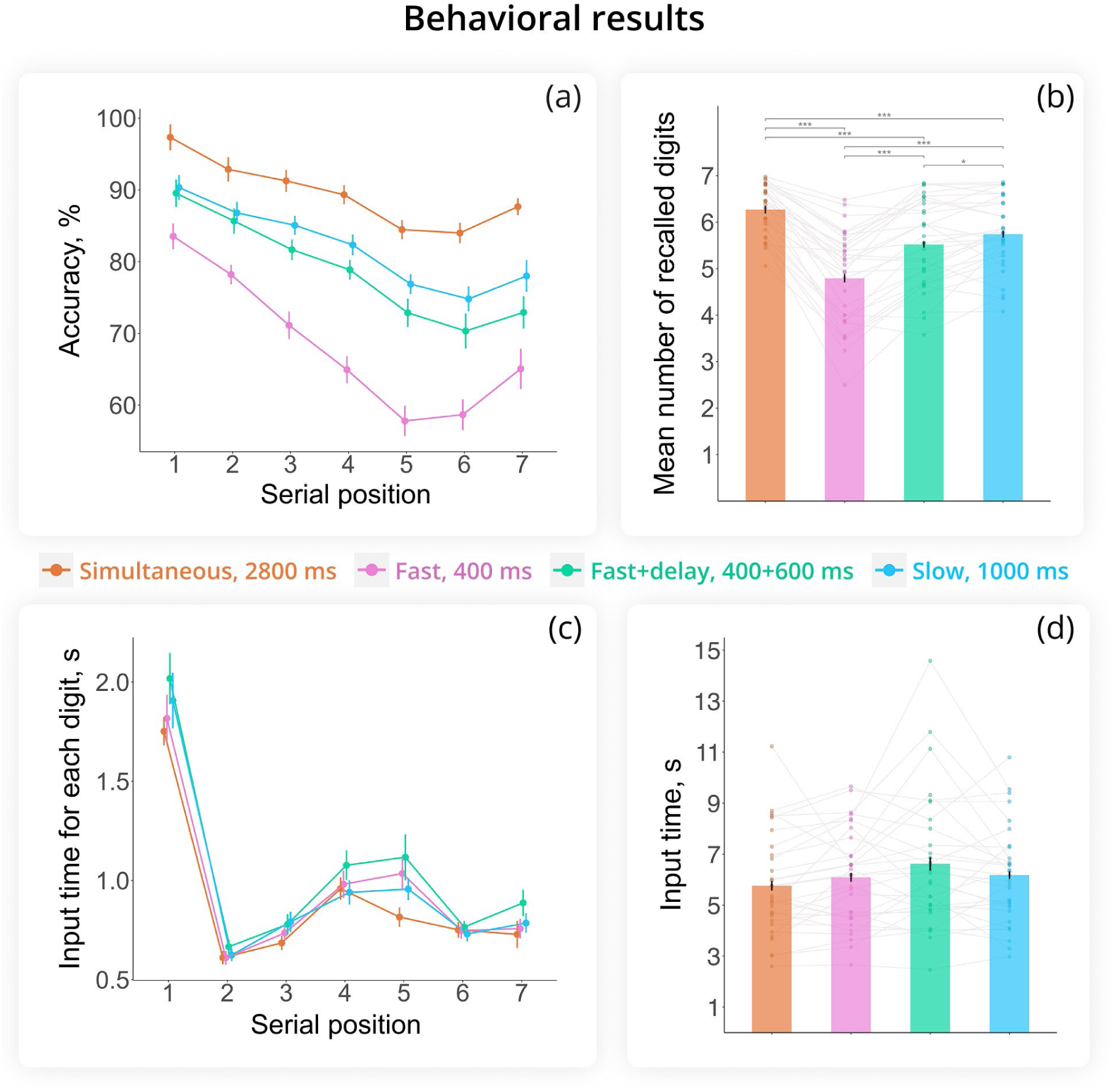
Behavioral performance. (a) Mean recall accuracy (%) as a function of presentation mode and the serial position of digits (WM load). (b) Mean number of digits recalled in the correct serial order for each condition (partial score). (c) Mean time taken to type each digit in the sequence across presentation modes. (d) Mean total input time, measured from the first keypress to the final ‘enter’ keypress. Error bars indicate the standard error of the mean (SEM).

Accuracy across serial positions varied as a function of Presentation mode (Serial position × Presentation mode interaction: F(7.69, 223.08) = 5.14, p < .001, *η_p_²* = .15). As shown in Figure 2a, the first digits in the sequence were recalled more accurately than subsequent digits across all modes (p < .01 for the first digit vs. all others), demonstrating a primacy effect. Accuracy generally decreased as WM load increased (Serial position: F(1.65, 47.95) = 28.54, p < .001, *η_p_²* = .50). Additionally, a recency effect was observed only in the Fast (p = 0.005) and Simultaneous (p = 0.02) modes (i.e., better performance for the 7th than 6th digit). Pairwise comparisons between presentation modes were significant for each serial position, except for positions 1 (p = 0.56), 2 (p = 0.45), 5 (p = 0.06), and 6 (p = 0.051) in the Slow vs. Fast+delay modes.

To estimate the duration of memory retrieval, we measured the time participants took to type their answers, from the first keypress to pressing ‘enter’. Participants spent less time typing in the Simultaneous mode (5.68 s) compared to the Fast+delay mode (6.60 s) (Presentation mode: F(2.28, 66.15) = 3.16, p = .043, *η_p_²* = .10; see Figure 2d). However, post hoc t-tests indicated that no pairwise comparisons survived adjustment for multiple comparisons.

Input time varied by Serial position (F(2.05, 59.32) = 69.88, p < .001, *η_p_²* = .706; Figure 2c). Participants took the longest time to enter the first digit, the shortest time to enter the second digit, and their input time slowed down in the middle of the sequence – a signature of chunking formation (Adam et al., 2024). The interaction between Serial position and Presentation mode was not significant (F(6.58, 190.78) = 1.33, p = .239, *η_p_²* = .04), suggesting similar retrieval patterns across presentation modes.

Because there were more trials for the first digit than for the other digits (due to the inclusion of the trials with attempted corrections), we analyzed the reaction time for the first digit separately. The mean reaction time did not differ by Presentation mode (F(2.57, 74.45) = 2.34, p = .090, *η_p_²* = .075).

### 3.2 Electroencephalography

### Alpha

#### Task period: encoding and retention

We found no significant difference in overall alpha power between presentation modes (F(2.49, 72.24) = 2.32, p = .093, *η_p_²* = .074). Generally, alpha activity was strongly suppressed during the encoding period (the time of digit presentation), increased above baseline during the first half of the retention period (0.6-3 s after retention onset), and gradually suppressed again during the second half (3-6 s) (Period: F(1.32, 38.36) = 26.36, p < .001, *η_p_²* = .48) (see Figure 2). This dynamics was influenced by Presentation mode (Period x Presentation mode: F(3.45, 100.07) = 5.71, p < .001, *η_p_²* = .16).

During encoding, alpha suppression was greater in Simultaneous (p = .01), Fast (p = .03) and Slow (p = .003) modes compared to Fast+Delay. In the first half of the retention period (0.6-3 s), alpha power increased in all modes except Fast+Delay (Figure 3b), where alpha went below baseline and was significantly weaker than in other presentation modes (p = .02 vs Simultaneous, p = .002 vs Fast, p = .01 vs Slow). In the second half of the retention period (3-6 s), alpha activity did not differ between presentation modes.

**Figure 3.**
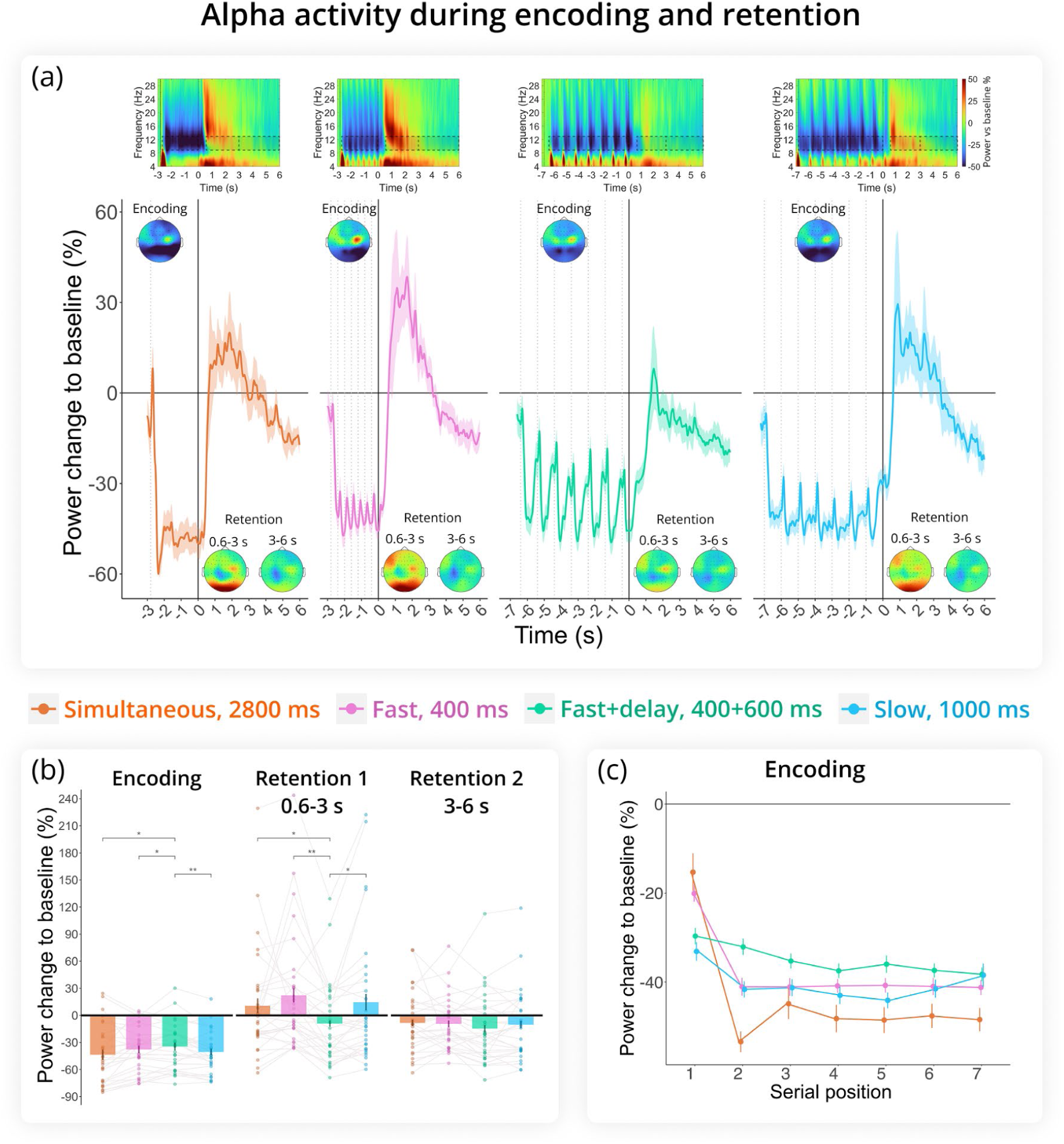
Alpha rhythm (9-13 Hz) results. (a) Change in alpha power relative to baseline across encoding and retention intervals, with 0 marking the onset of retention. Time-frequency maps at representative channels (PO7, PO3, POz, PO4, PO8, O1, Oz, O2) are shown at the top. Dashed boxes indicate time frequency regions of interest included in the analysis. Topograms illustrate alpha activity in the same cluster during Encoding, Retention 1, and Retention 2. Dashed lines indicate memory item presentation during encoding. (b) Change in alpha power relative to baseline averaged across the three task periods (Encoding, Retention 1, Retention 2). Fast+delay was the only condition that dropped below baseline during the first 3 s of retention. (c) Serial position effect during encoding. Alpha power was averaged over digit-presentation windows (400 ms for Fast, 1000 ms for Slow and Fast+delay, and seven 400 ms windows for Simultaneous). Error bars are SEM.

Further comparisons of sequential (all three combined) versus simultaneous presentation modes during the retention interval - also divided into two parts - revealed no significant effect of presentation mode (F(1, 29) = 0.15, p = .704, *η_p_²* < .01).

#### Encoding: serial position effect

Next, we assessed the effect of Serial position on alpha power during encoding using linear regression analysis. For this analysis, we averaged alpha activity over the time between each digit onset for the sequential conditions: 1000 ms in Fast+delay and Slow modes, and 400 ms in Fast mode. In the Simultaneous mode, we averaged alpha activity over seven 400 ms time windows to illustrate its time course (see Figure 3c). An omnibus ANOVA was conducted to assess the significance of the categorical variable Presentation mode.

Although the ANOVA indicated an overall effect of Presentation mode on the mean alpha power during encoding (Presentation mode: F = 3.81, p = .01), pairwise comparisons did not reveal any differences between conditions. Additionally, the amount of suppression increased with each digit to be encoded (Serial position: F = 16.01, p < .001). The interaction between Serial position and Presentation mode was not significant (F = 1.46, p = .22).

Because there was an exceptionally strong change in alpha at the onset of the first digit in the sequence, the Serial position effect might have been highly significant due to this effect alone. When only digits 2 to 7 were included in the analysis, the overall alpha suppression distribution remained similar to the previous analysis, with Simultaneous mode eliciting stronger alpha suppression than Fast+delay mode (Presentation mode: F = 6.65, p < .001; Simultaneous vs Fast+delay: t = 2.42, p = .02). However, the factor Serial position lost significance (F = 0.005, p = .945), indicating that alpha power did not parametrically decrease with increasing WM load during the encoding period. Moreover, the Serial position effect was similar across conditions (Serial position × Presentation mode: F = 0.35, p = .788).

### Theta

#### Task period: encoding and retention

As depicted in Figure 4a, theta activity increased during encoding, peaked at the beginning of the retention interval, and then gradually decreased until the end of retention. However, an ANOVA revealed that theta activity did not differ significantly between task periods (F(1.15, 33.25) = 3.15, p = .080, *η_p_²* = .10) and was not significantly affected by presentation mode (F(2.26, 65.49) = 0.85, p = .443, *η_p_²* = .03). Additionally, the interaction between Period and Presentation mode was not significant (F(1.96, 56.73) = 0.18, p = .832, *η_p_²* < .01).

**Figure 4.**
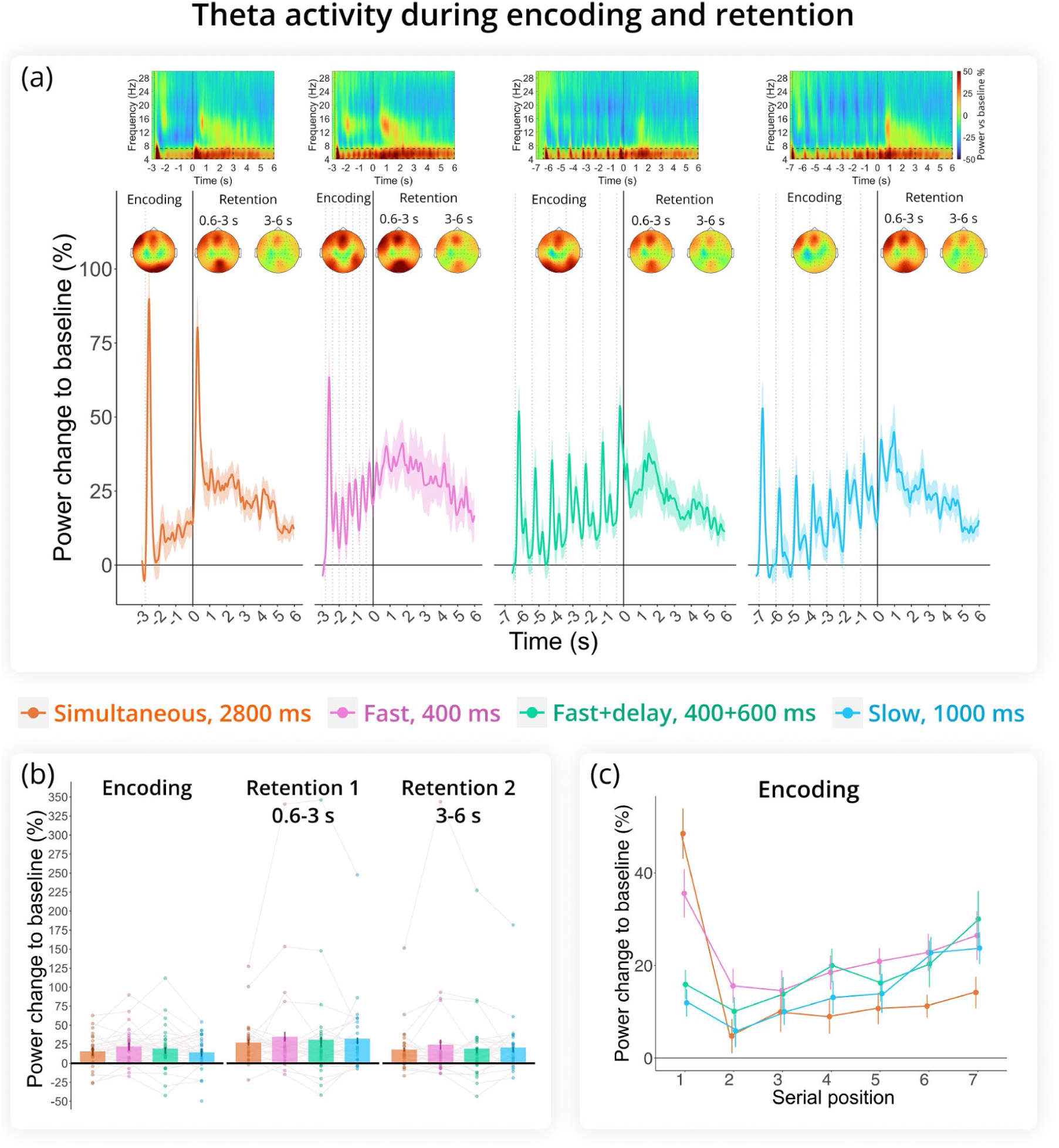
Theta rhythm (4-7 Hz) results. (a) Change in theta power relative to baseline across encoding and retention intervals, with 0 marking the onset of retention. Time-frequency maps at Fz appear at the top. Dashed boxes indicate time frequency regions of interest included in the analysis. Topograms illustrate theta activity in the same cluster during Encoding, Retention 1, and Retention 2. Dashed lines indicate memory item presentation during encoding. (b) Change in theta power relative to baseline averaged across the three task periods (Encoding, Retention 1, Retention 2). (c) Serial position effect during encoding. Theta power was averaged over digit-presentation windows (400 ms for Fast, 1000 ms for Slow and Fast+delay, and seven 400 ms windows for Simultaneous). Error bars are SEM.

#### Encoding: serial position effect

Considering encoding period separately, linear regression analysis showed that mean theta power was consistent across conditions (ANOVA omnibus test for the effect of Presentation mode: F = 2.579, p = .052) and did not change with increasing cognitive load (Serial position: F = 0.520, p = .47). Within conditions, there was a change in theta activity with the presentation of each digit for encoding (Serial position x Presentation mode: F = 6.587, p < .001) such that in the Simultaneous and Fast conditions there was a tendency for theta to decrease with load, whereas in the Slow and Fast+delay conditions there was an upward trend.

As with the alpha rhythm, there was a strong surge in theta activity at the presentation of the first digit in the sequence which might reflect some process different from WM. Therefore, we repeated the analysis without the first digit. Although the general effect of Presentation mode was significant (F = 3.805, p = .01), no pairwise comparisons between conditions attained the significance level. Importantly, theta showed an upward slope with increasing load (Serial position: F = 16.78, p < .001), with the trend for theta growth being similar in all conditions (Serial position x Presentation mode: F = 0.55, p = .647).

### Beta

#### Task period: encoding and retention

Overall, presentation mode did not significantly affect beta power dynamics in the task (F(2.71, 78.48) = 2.31, p = .089, *η_p_²* = .074). Beta power decreased during the encoding period, remained below baseline during first half of the retention, and continued to decrease in the second half (Period: F(1.52, 44.17) = 24.33, p < .001, *η_p_²* = .46; all pairwise comparisons yielded p < .005; see Figure 5b). There was a significant interaction between Period and Presentation mode (F(4.57, 132.61) = 4.80, p < .001, *η_p_²* = .14). This interaction was driven by significantly stronger beta suppression during encoding in the Slow mode compared to the Fast (p = 0.005) and Simultaneous modes (p = 0.03). Although this pattern persisted across all task periods, it was only statistically significant during Encoding (Presentation mode: F(2.96, 85.81) = 4.88, p = 0.004, *η_p_²* = .14).

**Figure 5.**
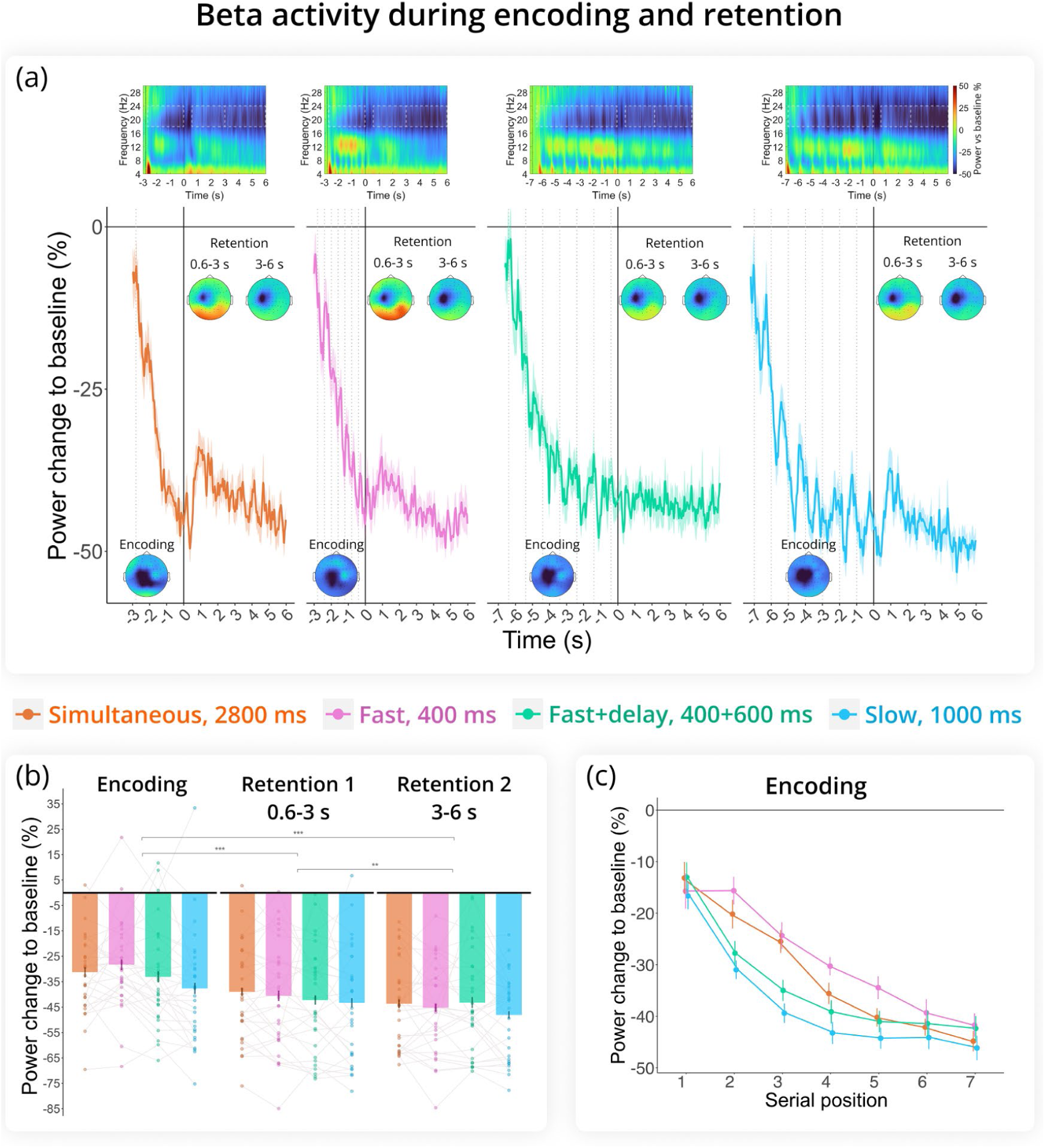
Beta rhythm (18-24 Hz) results. (a) Change in beta power relative to baseline across encoding and retention intervals, with 0 marking the onset of retention. Time-frequency maps at C3 are shown at the top. Dashed boxes indicate time frequency regions of interest included in the analysis. Topograms illustrate beta activity in the same cluster during Encoding, Retention 1, and Retention 2. Dashed lines indicate memory item presentation during encoding. (b) Change in beta power relative to baseline averaged across the three task periods (Encoding, Retention 1, Retention 2). (c) Serial position effect during encoding. Beta power was averaged over digit-presentation windows (400 ms for Fast, 1000 ms for Slow and Fast+delay, and seven 400 ms windows for Simultaneous). Error bars are SEM.

#### Encoding: serial position effect

Linear regression analysis of the encoding period revealed that the Simultaneous (t = 2.408, p = 0.016) and Fast (t = 2.475, p = 0.014) presentation modes exhibited overall weaker suppression compared to the Slow mode (Presentation mode: F = 7.07, p < .001). Suppression also increased gradually with WM load producing a strong main effect of Serial position (F = 171.11, p < .001; Figure 5c). However, the rate of decrease in beta power was similar across all conditions (Serial position × Presentation mode: F = 0.60, p = .61).

### Combined single-trial accuracy-EEG analysis

During encoding, alpha (b = 0.218, SE = 0.088, t = 2.47, p = 0.013) and theta (b = 0.213, SE = 0.079, t = 2.70, p = 0.007) power predicted performance, with stronger theta enhancement and deeper alpha suppression associated with more digits recalled. Beta effects during encoding were not significant (b = 0.035, SE = 0.107, t = 0.32, p = 0.746). Both alpha and theta results remained significant after removing the first 400 ms of encoding. No effects emerged in the retention period (parts 1 and 2).

## 4 Discussion

Our primary objective was to determine whether stimulus presentation mode influences alpha activity in a WM task, particularly with respect to the direction of alpha changes in the retention interval. The findings revealed that alpha activity is indeed sensitive to presentation mode in both encoding and retention periods of the task. Participants achieved their highest recall accuracy under the simultaneous presentation mode, outperforming all sequential conditions. Despite this behavioral advantage, alpha levels during retention did not differ between simultaneous and sequential modes. In contrast, alpha power showed a stark difference between the Slow (1000 ms presentation) and Fast+delay (400 ms presentation interspersed with 600 ms breaks) sequential modes. Specifically, on average, alpha power was below baseline in the Fast+delay condition but well above baseline in the Slow condition - despite comparable behavioral accuracy. During encoding, alpha suppression was strongest in the Simultaneous mode and weakest in the Fast+delay mode.

Whereas alpha power did not significantly change with increasing WM load during the encoding period in any presentation mode, frontal midline theta power uniformly increased with WM load regardless of presentation mode during encoding and remained independent of presentation mode during the retention period. Meanwhile, beta power localized to the left motor cortex remained consistently below baseline and became progressively more suppressed with each additional digit memorized. Although this suppression was stronger in the Slow mode than in modes with a shorter SOA, it was independent of the presentation mode during the retention period.

Participants recalled digit sets most accurately when the digits were presented simultaneously, even though the overall encoding time for Simultaneous mode (2800 ms) was shorter than that for Slow mode (7000 ms). This aligns with evidence that simultaneous presentation of verbal material, compared with sequential presentation of the same number of items without blank intervals, yields better performance (Ordonez Magro et al., 2022; Ricker & Cowan, 2014). The presence of primacy and recency, and the accuracy distribution within Simultaneous, indicate that participants still encoded the list in a left to right order, so the advantage likely reflects more efficient encoding while preserving the presented left to right recall order. Parallel reading and stronger item-position binding (Bharti et al., 2020; Ordonez Magro et al., 2022) as well as the facilitation of chunking may both contribute to this improvement in performance (Cowan et al., 2012). Simultaneous presentation is not uniformly advantageous across domains. In visual tasks the benefit is often absent when overall exposure is matched across formats (Chen & Kuo, 2025; Johnson et al., 2009). Our sequential conditions illustrate when the gap narrows. Slow improved accuracy relative to Fast by increasing per item exposure, and Fast+delay improved accuracy by inserting brief pauses (Mızrak & Oberauer, 2021; Ricker & Cowan, 2014). These patterns suggest that for verbal material the most effective approach is to present all items at once when possible, or if a sequential format is required, to provide sufficient exposure or inter-item pauses.

Despite substantial differences in behavioral performance, we found no significant differences in alpha activity when comparing simultaneous and sequential presentation modes. This result contrasts with the findings of Okuhata et al. (2013), who reported lower alpha power in the simultaneous than in the sequential mode. It is important to note that, in their study, digits were displayed around a fixation cross in the simultaneous condition, potentially altering verbal processing accessibility and promoting visual strategies. One attempt to reconcile conflicting results regarding alpha modulation in WM tasks comes from the proposal that posterior alpha power tends to increase with WM load in tasks relying on verbal strategies but decreases when the task emphasizes maintaining visual identity (Chen et al., 2022). According to this view, alpha power decreases as WM load grows if the same neural circuits support both encoding and retention, whereas it increases if encoding and retention engage different circuits. Although this framework is compelling, numerous studies involving matching-to-sample tasks involving verbal material have reported alpha increases (Pavlov & Kotchoubey, 2022), while our previous study using a digit span paradigm showed alpha decreases (Kosachenko et al., 2023). Our review likewise indicates that alpha increases in roughly half of the reported cases for visual WM tasks (Pavlov & Kotchoubey, 2022). In the current study, simply adding a brief delay between memory items pushed average parieto-occipital alpha power below baseline level. This shift cannot be explained by a transition from verbal to visual encoding. Thus, attributing alpha differences in verbal WM tasks solely to the involvement of the visual cortex is not supported by both the existing literature and the current findings.

Longer encoding time in a sequential presentation mode - from 400 ms to 1000 ms - has previously been shown to improve memory accuracy (Oberauer, 2022). Additionally, our results indicate that providing extra free time between memory items (as in the Fast+delay compared to Fast mode) also enhances performance. This finding aligns with the evidence that increased free time between memory items during encoding boosts accuracy (Mızrak & Oberauer, 2021). This boosting effect can be explained by the process of WM consolidation (Oberauer, 2022; Ricker & Cowan, 2014; Ricker et al., 2018), during which fleeting perceptions are converted into more stable WM representations, and more consolidation time reduces the rate of forgetting. When consolidation time is insufficient - less than approximately 500-1000 ms (Ricker et al., 2018) - memory performance declines compared with conditions with adequate time. In our study, the weaker performance in the 400-ms condition can be attributed to this lack of consolidation time. Previous studies typically fixed the duration of encoding (e.g., 250 ms in Ricker & Cowan (2014)) while varying only the free time between memory items. By examining both effects of free time intervals and encoding duration, our study provides further evidence for the beneficial role of inter-item free time. Moreover, our data indicate that an increased encoding duration can replace a shorter encoding period plus additional free time, since both presentation modes yield similar levels of accuracy. However, as shown by EEG data, the neural processes underlying similar behavioral performance may differ.

Despite comparable recall accuracy, the Slow and Fast+delay presentation modes exhibited striking differences in alpha power during encoding and retention. In the Slow mode, alpha increased only slightly during each digit presentation but rose sharply during retention. By contrast, in Fast+delay, alpha increased more robustly during digit presentation, then showed a markedly weaker enhancement once retention began. Simply attributing this pattern to the greater number of stimulus changes in Fast+delay is insufficient, since each stimulus would normally be expected to elicit alpha suppression (Pfurtscheller & Aranibar, 1977), with cumulative suppression lasting longer when more stimuli are presented. A plausible account is that the repeated fixation cross intervals blur the boundary between encoding and retention, delaying the shift into retention. Consistent with this, the early-retention alpha increase in Fast+delay lagged Slow by about 600 ms, matching the inter-item pause, despite identical visual input at retention onset. Participants knew that each list contained seven digits, so the transition was predictable, yet the visual regularity could still delay the early retention response. This can explain the delayed onset but not the reduced magnitude of the alpha increase in Fast+delay, and a simple sensory rebound account is also unlikely given the identical visual input at retention onset across modes. A further possibility is that the fixation cross promoted a shift of attention from external input to internal representations, enabling visual cortical disengagement that may coincide with working memory consolidation.

Additional consolidation time can bolster resistance to attention-demanding interference (Barrouillet et al., 2013). This may explain why alpha surges appear after each memory item, yet there is a smaller alpha increase during retention in Fast+delay - interference resistance may have been deferred to the moments immediately after each item’s encoding. The alpha surges can indicate the inhibition of visual sensory areas coinciding with WM consolidation in the intervals between active encoding, paralleling long-term memory (LTM) consolidation processes that occur between active learning states (Dudai et al., 2015). This inhibition during the retention interval may only be necessary while the memory traces are fragile and not yet consolidated. Indeed, further extending encoding and between-item delay durations (to 1 s each) has been shown to result in effectively eliminating the alpha increase typically observed at the transition from encoding to retention (Kosachenko et al., 2023). This suggests that the alpha-related mechanisms supporting WM function are no longer essential once memory traces have had sufficient time to stabilize (possibly, between 1 and 2 seconds).

Although encoding and consolidation appear to compete for cognitive resources, they are not mutually exclusive. Encoding may be prioritized when feasible, perhaps because it is less resource-intensive than consolidation. This flexible allocation of limited cognitive resources - signaled by alpha - between encoding and retention did not substantially affect overall performance but highlights a qualitative shift in how memory is formed. Transitioning from passive encoding to a more consolidation-centric strategy affects processing and retaining information, suggesting distinct underlying mechanisms in the Slow vs. Fast+delay modes. However, our inference that consolidation processes drive alpha power dynamics is based on limited evidence and must be rigorously tested in future studies.

Consolidation processes likely contribute not only to stabilizing content of WM but also to transferring it to LTM. Verbal WM’s limited capacity often leverages LTM to handle higher loads (Bartsch & Oberauer, 2023; Cowan et al., 2012). Cotton & Ricker (2021) showed that consolidation in WM, rather than mere attention, is crucial for enhancing LTM performance. Studies also indicate that providing additional free time immediately post-item presentation can bolster subsequent LTM retention (Hartshorne & Makovski, 2019; Jarjat et al., 2018, 2020; Souza & Oberauer, 2017). Although speculative, our findings suggest that the Fast+delay condition could foster a setting conducive to more active LTM engagement, evidenced by the longest retrieval time. This extended retrieval window may reflect deeper consolidation, producing robust yet less immediately accessible traces. The deeper consolidation may also make mental rehearsal during retention period more resource intensive and thus more alpha suppressing if alpha is only a representation of non-specific cortical activation. However, the retrieval effect was weak and did not survive correction for multiple comparisons. Nevertheless, weaker alpha enhancement during retention has been linked to greater LTM involvement. For instance, Kasanov et al. (2024) reported reduced alpha enhancement when WM contents were manipulated alphabetically (which requires semantic LTM) vs. backward reordering (which does not). Similarly, Berger et al. (2014) and Pavlov & Kotchoubey (2021) observed decreased alpha when LTM access was required, as compared with simple retention. These observations align with the idea that alpha progressively relaxes sensory cortical inhibition when LTM must integrate with WM (Klimesch, 2012; Klimesch et al., 2007). Thus, our data further reinforce the link between alpha enhancement strength and LTM engagement during retention.

Within modes, trials with deeper alpha suppression during encoding were recalled better. This supports the view that performance depends on effective sensory engagement during encoding, even in Fast+delay where inter-item pauses raise mean encoding alpha via rebounds. Importantly, retention alpha was unrelated to accuracy. To draw a stronger inference about the role of retention alpha in working memory, future work should rule out that the alpha increase reflects perceptual load rather than working memory per se. Designs that vary set size across trials and include a matched visual control without memory demands can help separate sensory driven from working memory specific components.

Theta rhythm increased as expected during encoding, scaling with rising WM load. In contrast to alpha, the presentation format had no effect on theta - whether items were shown every 400 ms, 1000 ms, or all at once for 2800 ms, the theta loading function remained unchanged. Typically, theta rhythm is associated with executive control networks within WM (Kosachenko et al., 2023; Pavlov & Kotchoubey, 2020) or beyond (Cavanagh & Frank, 2014). Our results suggest that the executive control processes reflected in frontal midline theta activity during encoding and retention do not depend on presentation mode - simultaneous or sequential - or on the pace of sequential presentation. The observed theta pattern fits an executive control account that is engaged when items are added and ordered, which helps explain why the 400 ms condition did not elevate theta during encoding despite higher sensory difficulty. At the single trial level, stronger theta during encoding predicted better recall accuracy, consistent with theta indexing control processes that order and stabilize items as they are added to memory.

Centrally localized beta activity remained below baseline throughout the trial and became more suppressed with each digit presentation during the encoding period. By around 4 s, the longer SOA conditions reached the same level of beta suppression and maintained it, while Fast and Simultaneous modes displayed a steeper loading function before converging around the sixth digit. During the retention period, beta activity remained at similar levels across all presentation modes. Beta suppression in the sensorimotor cortex contralateral to the required response hand has been associated with response planning (Boettcher et al., 2021; Schneider et al., 2017; van Wijk et al., 2009) and response sequence learning (Zhuang et al., 1997). In our task, response planning tracked by beta suppression during encoding was facilitated by the straightforward mapping of digits on the numpad, as well as the requirement to fully reproduce the sequence by typing. We propose that each new digit in the sequence initiates a parallel encoding process that forms both memory representations in the sensory cortex and response planning. Differences in beta activity across presentation modes in our study are likely driven by the pace of item presentation, which can deprioritize and/or delay motor planning so that it engages only if time permits - either between the presentation of memory items or after all have been encoded by the sensory mechanism. Previous studies have also shown concurrent response planning and memory encoding/retention of visual WM (Boettcher et al., 2021; Nasrawi et al., 2025; Schneider et al., 2017). We showed that the signature of motor planning extends to verbal WM encoding.

Consistent with a motor preparation account, beta did not predict recall accuracy. Errors in this task arise from item loss or order swaps rather than motor execution failure, so the accuracy metric depends on the integrity of mnemonic content, whereas central beta indexes readiness to type the response, whether correct or not. Prior work links central beta to action preparation and sequence execution rather than mnemonic quality (Nasrawi et al., 2023, 2025).

## 4.1 Conclusions

Our results show that alpha, theta, and beta track complementary facets of verbal working memory that are shaped by the timing of item presentation. Alpha power can increase or decrease during verbal working memory retention, depending not so much on presentation mode but, rather, on how much “breathing room” is allowed between items. Specifically, adding brief pauses after each item (Fast+delay) drove alpha below baseline during retention, whereas both faster and slower sequential presentations without breaks - and the behaviorally superior simultaneous presentation - resulted in elevated alpha. Single trial models linked deeper alpha suppression during encoding, not retention alpha, to better recall. Our results indicate that alpha may reflect more than sensory engagement during encoding or the suppression of irrelevant sensory areas, it also appears to capture consolidation processes. Theta increased with memory load across all modes during encoding and predicted recall on a trial by trial basis, consistent with an executive control signal that orders and stabilizes items as they are added. Beta over central motor cortex decreased with load during encoding and reflected preparation for serial report, yet it did not predict accuracy, in line with its role in action planning rather than mnemonic fidelity.

## Author contributions

Alexandra I. Kosachenko: Conceptualization; data curation; formal analysis; investigation; methodology; visualization; writing – original draft; writing – review and editing.

Danil I. Syttykov: Investigation; methodology.

Alexander I. Kotyusov: Investigation; methodology.

Dmitry A. Tarasov: Investigation.

Dauren Kasanov: Investigation.

Sergey Malykh: Writing – review and editing.

Boris Kotchoubey: Writing – review and editing.

Yuri G. Pavlov: Conceptualization; data curation; formal analysis; funding acquisition; methodology; project administration; supervision; writing – original draft; writing – review and editing.

## Acknowledgments

We thank Niko Busch and Agatha Lenartowicz for their valuable comments on an earlier draft of the manuscript.

## Data Availability Statement

Raw EEG data and stimulus presentation code are publicly available on Openneuro (https://openneuro.org/datasets/ds006848). Analysis code is available on OSF (https://osf.io/8n4a6/).

## Abbreviations

ANOVA: Analysis of Variance
AMICA: Adaptive Mixture Independent Component Analysis
BH: Benjamini-Hochberg procedure
EEG: Electroencephalography
LTM: Long-Term Memory
RM: Repeated Measures
ROI: Region of Interest
SD: Standard Deviation
SEM: Standard Error of the Mean
SOA: Stimulus Onset Asynchrony
WM: Working Memory

## Supplementary Results

Cluster-based permutation tests were run separately for each frequency band and task period, comparing presentation modes pairwise. The analysis was conducted in Fieldtrip toolbox (Oostenveld et al., 2011) with 5,000 permutations, a cluster alpha of 0.05, and a minimum cluster size of 2 neighboring channels.

**Figure S1.**
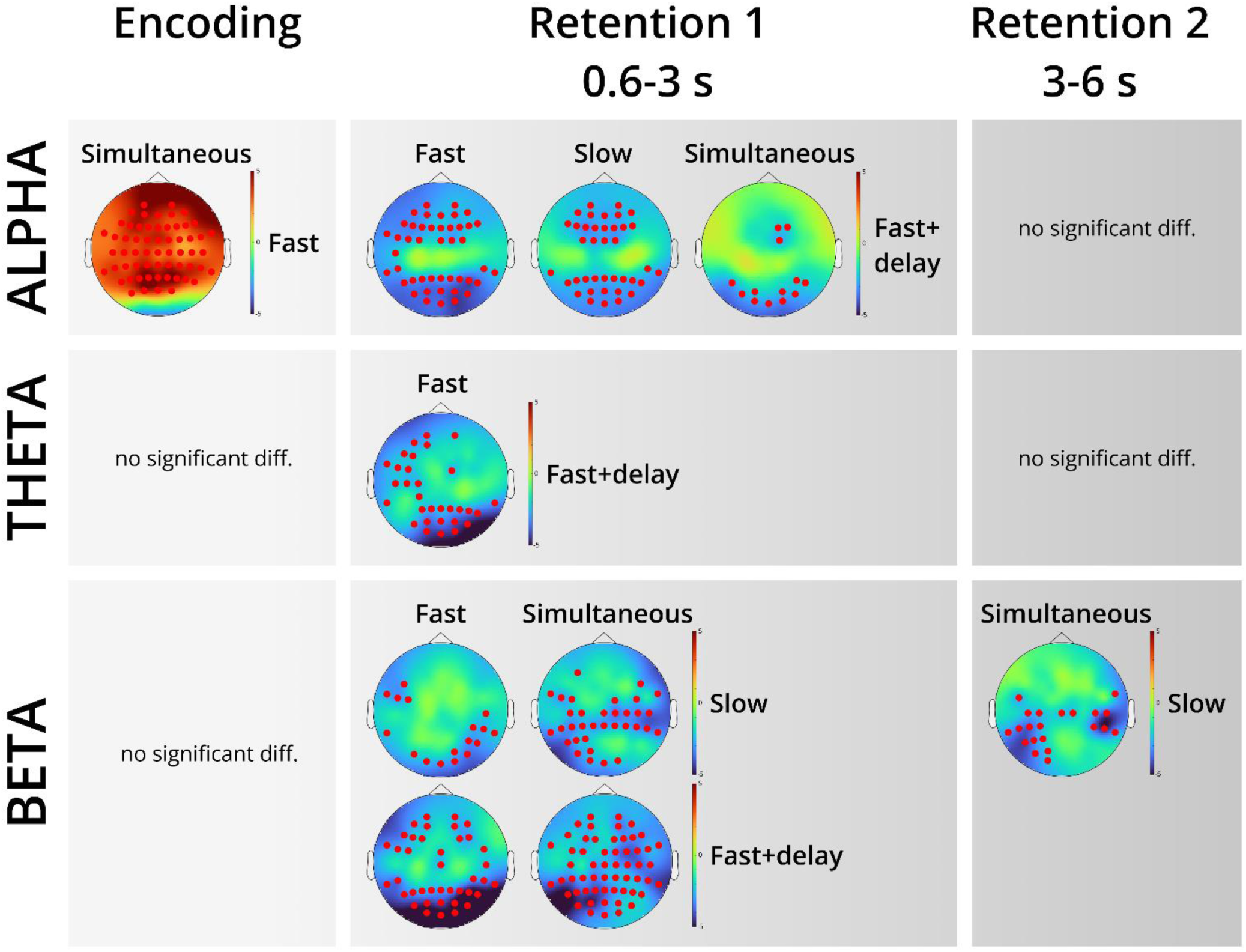
Results of cluster-based permutation tests across all electrodes for the alpha (9-13 Hz), theta (4-7 Hz), and beta (18-24 Hz) frequency bands during (1) Encoding, (2) Retention 1 (0.6-3 s), and (3) Retention 2 (3-6 s). The color scale represents t-values, with positive values indicating stronger activation in the presentation mode listed on the right compared with the condition listed at the top (e.g., Fast vs. Simultaneous for alpha during encoding). Red dots indicate electrodes showing significant differences between the compared conditions. Only significant effects are displayed.

## References

Adam, K. C. S., Zhao, C., & Vogel, E. K. (2024). Behavioral signatures of the rapid recruitment of long-term memory to overcome working memory capacity limits. Memory & Cognition, 52(8), 1816–1832. 10.3758/s13421-024-01566-z

Bartsch, L. M., & Oberauer, K. (2023). The contribution of episodic long-term memory to working memory for bindings. Cognition, 231, 105330. 10.1016/j.cognition.2022.105330

Benjamini, Y., & Hochberg, Y. (1995). Controlling the False Discovery Rate: A Practical and Powerful Approach to Multiple Testing. Journal of the Royal Statistical Society: Series B (Methodological), 57(1), 289–300. 10.1111/j.2517-6161.1995.tb02031.x

Berger, B., Omer, S., Minarik, T., Sterr, A., & Sauseng, P. (2014). Interacting Memory Systems—Does EEG Alpha Activity Respond to Semantic Long-Term Memory Access in a Working Memory Task? Biology, 4(1), 1–16. 10.3390/biology4010001

Bharti, A. K., Yadav, S. K., & Jaswal, S. (2020). Feature Binding of Sequentially Presented Stimuli in Visual Working Memory. Frontiers in Psychology, 11. 10.3389/fpsyg.2020.00033

Boettcher, S. E. P., Gresch, D., Nobre, A. C., & van Ede, F. (2021). Output planning at the input stage in visual working memory. Science Advances, 7(13), eabe8212. 10.1126/sciadv.abe8212

Bowman, H., Brooks, J. L., Hajilou, O., Zoumpoulaki, A., & Litvak, V. (2020). Breaking the circularity in circular analyses: Simulations and formal treatment of the flattened average approach. PLOS Computational Biology, 16(11), e1008286. 10.1371/journal.pcbi.1008286

Brickwedde, M., Limachya, R., Markiewicz, R., Sutton, E., Postzich, C., Shapiro, K., … Mazaheri, A. (2025). Cross-modal interaction of Alpha Activity does not reflect inhibition of early sensory processing: A frequency tagging study using EEG and MEG. eLife, 14. 10.7554/eLife.106050.3

Cavanagh, J. F., & Frank, M. J. (2014). Frontal theta as a mechanism for cognitive control. Trends in Cognitive Sciences, 18(8), 414–421. 10.1016/j.tics.2014.04.012

Chen, Y.-T., & Kuo, B.-C. (2025). Simultaneous and Sequential Presentations Differentially Modulate the Temporal Dynamics of Working Memory Processes. Journal of Cognitive Neuroscience, 1–15. 10.1162/JOCN.a.2399

Chen, Y.-T., Van Ede, F., & Kuo, B.-C. (2022). Alpha Oscillations Track Content-Specific Working Memory Capacity. The Journal of Neuroscience, 42(38), 7285–7293. 10.1523/JNEUROSCI.2296-21.2022

Cotton, K., & Ricker, T. J. (2021). Working memory consolidation improves long-term memory recognition. *Journal of Experimental Psychology: Learning*, Memory, and Cognition, 47(2), 208–219. 10.1037/xlm0000954

Cowan, N., Rouder, J. N., Blume, C. L., & Saults, J. S. (2012). Models of verbal working memory capacity: What does it take to make them work? Psychological Review, 119(3), 480–499. 10.1037/a0027791

Delorme, A., & Makeig, S. (2004). EEGLAB: An open source toolbox for analysis of single-trial EEG dynamics including independent component analysis. Journal of Neuroscience Methods, 134(1), 9–21. 10.1016/j.jneumeth.2003.10.009

Erickson, M. A., Albrecht, M. A., Robinson, B., Luck, S. J., & Gold, J. M. (2017). Impaired suppression of delay-period alpha and beta is associated with impaired working memory in schizophrenia. Biological Psychiatry. Cognitive Neuroscience and Neuroimaging, 2(3), 272–279. https://doi.org/10/gf27qt

Gathercole, S. E. (2012). Working Memory and Language. In M. G. Gaskell (Ed.), The Oxford Handbook of Psycholinguistics (1st ed., pp. 757–770). 10.1093/oxfordhb/9780198568971.013.0046

Gignac, G. E. (2014). Fluid intelligence shares closer to 60% of its variance with working memory capacity and is a better indicator of general intelligence. Intelligence, 47, 122–133. 10.1016/j.intell.2014.09.004

Hartshorne, J. K., & Makovski, T. (2019). The effect of working memory maintenance on long-term memory. Memory & Cognition, 47(4), 749–763. 10.3758/s13421-019-00908-6

Hitch, G. J., Allen, R. J., & Baddeley, A. D. (2024). The multicomponent model of working memory fifty years on. Quarterly Journal of Experimental Psychology, 17470218241290909. 10.1177/17470218241290909

Jarjat, G., Hoareau, V., Plancher, G., Hot, P., Lemaire, B., & Portrat, S. (2018). What makes working memory traces stable over time? Annals of the New York Academy of Sciences, 1424(1), 149–160. 10.1111/nyas.13668

Jarjat, G., Plancher, G., & Portrat, S. (2020). Core mechanisms underlying the long-term stability of working memory traces still work in aging: L’Année Psychologique, Vol. 120(2), 203–229. 10.3917/anpsy1.202.0203

Jensen, O. (2024). Distractor inhibition by alpha oscillations is controlled by an indirect mechanism governed by goal-relevant information. Communications Psychology, 2(1), 36. 10.1038/s44271-024-00081-w

Jensen, O., Gelfand, J., Kounios, J., & Lisman, J. E. (2002a). Oscillations in the alpha band (9-12 Hz) increase with memory load during retention in a short-term memory task. Cerebral Cortex, 12(8), 877–882. 10.1093/cercor/12.8.877

Jensen, O., Gelfand, J., Kounios, J., & Lisman, J. E. (2002b). Oscillations in the alpha band (9-12 Hz) increase with memory load during retention in a short-term memory task. Cerebral Cortex, 12(8), 877–882. 10.1093/cercor/12.8.877

Jensen, O., & Mazaheri, A. (2010). Shaping Functional Architecture by Oscillatory Alpha Activity: Gating by Inhibition. Frontiers in Human Neuroscience, 4. 10.3389/fnhum.2010.00186

Johnson, J. S., Spencer, J. P., Luck, S. J., & Schöner, G. (2009). A Dynamic Neural Field Model of Visual Working Memory and Change Detection. Psychological Science, 20(5), 568–577. 10.1111/j.1467-9280.2009.02329.x

Kasanov, D., Dorogina, O., Mushtaq, F., & Pavlov, Y. G. (2024, March 20). Theta transcranial alternating current stimulation is not effective in improving working memory performance. 10.1101/2024.03.20.585954

Klimesch, W. (2012). α-band oscillations, attention, and controlled access to stored information. Trends in Cognitive Sciences, 16(12), 606–617. 10.1016/j.tics.2012.10.007

Klimesch, W., Sauseng, P., & Hanslmayr, S. (2007). EEG alpha oscillations: The inhibition–timing hypothesis. Brain Research Reviews, 53(1), 63–88. 10.1016/j.brainresrev.2006.06.003

Kosachenko, A. I., Kasanov, D., Kotyusov, A. I., & Pavlov, Y. G. (2023). EEG and pupillometric signatures of working memory overload. Psychophysiology, 60(6), e14275. 10.1111/psyp.14275

Liljefors, J., Almeida, R., Rane, G., Lundström, J. N., Herman, P., & Lundqvist, M. (2024). Distinct functions for beta and alpha bursts in gating of human working memory. Nature Communications, 15(1), 8950. 10.1038/s41467-024-53257-7

Mızrak, E., & Oberauer, K. (2021). What Is Time Good for in Working Memory? Psychological Science, 32(8), 1325–1337. 10.1177/0956797621996659

Nasrawi, E., Boettcher, S. E. P., & Ede, F. van. (2023). Prospection of Potential Actions during Visual Working Memory Starts Early, Is Flexible, and Predicts Behavior. Journal of Neuroscience, 43(49), 8515–8524. 10.1523/JNEUROSCI.0709-23.2023

Nasrawi, R., Mautner-Rohde, M., & van Ede, F. (2025). Memory load influences our preparedness to act on visual representations in working memory without affecting their accessibility. Progress in Neurobiology, 245, 102717. 10.1016/j.pneurobio.2025.102717

Oberauer, K. (2022). When does working memory get better with longer time? Journal of Experimental Psychology: Learning, Memory, and Cognition, 48(12), 1754–1774. 10.1037/xlm0001199

Okuhata, S., Kusanagi, T., & Kobayashi, T. (2013). Parietal EEG alpha suppression time of memory retrieval reflects memory load while the alpha power of memory maintenance is a composite of the visual process according to simultaneous and successive Sternberg memory tasks. Neuroscience Letters, 555, 79–84. 10.1016/j.neulet.2013.09.010

Onton, J., Delorme, A., & Makeig, S. (2005). Frontal midline EEG dynamics during working memory. NeuroImage, 27(2), 341–356. 10.1016/j.neuroimage.2005.04.014

Oostenveld, R., Fries, P., Maris, E., & Schoffelen, J.-M. (2011). FieldTrip: Open Source Software for Advanced Analysis of MEG, EEG, and Invasive Electrophysiological Data. Computational Intelligence and Neuroscience, 2011, 1–9. 10.1155/2011/156869

Ordonez Magro, L., Mirault, J., Grainger, J., & Majerus, S. (2022). Sequential versus simultaneous presentation of memoranda in verbal working memory: (How) does it matter? Memory & Cognition, 50(8), 1756–1771. 10.3758/s13421-022-01284-4

Palmer, J. A., Kreutz-Delgado, K., & Makeig, S. (2011). AMICA: An Adaptive Mixture of Independent Component Analyzers with Shared Components. 16.

Pavlov, Y. G., & Kotchoubey, B. (2020, May 3). The electrophysiological underpinnings of variation in verbal working memory capacity. 10.1101/2020.05.02.073825

Pavlov, Y. G., & Kotchoubey, B. (2021). Temporally distinct oscillatory codes of retention and manipulation of verbal working memory. European Journal of Neuroscience, 54(7), 6497–6511. 10.1111/ejn.15457

Pavlov, Y. G., & Kotchoubey, B. (2022). Oscillatory brain activity and maintenance of verbal and visual working memory: A systematic review. Psychophysiology, 59(5), e13735. 10.1111/psyp.13735

Peirce, J., Gray, J. R., Simpson, S., MacAskill, M., Höchenberger, R., Sogo, H., … Lindeløv, J. K. (2019). PsychoPy2: Experiments in behavior made easy. Behavior Research Methods, 51(1), 195–203. 10.3758/s13428-018-01193-y

Pfurtscheller, G., & Aranibar, A. (1977). Event-related cortical desynchronization detected by power measurements of scalp EEG. Electroencephalography and Clinical Neurophysiology, 42(6), 817–826. 10.1016/0013-4694(77)90235-8

Ratcliffe, O., Shapiro, K., & Staresina, B. P. (2022). Fronto-medial theta coordinates posterior maintenance of working memory content. Current Biology, 32(10), 2121–2129.e3. 10.1016/j.cub.2022.03.045

Ricker, T. J., & Cowan, N. (2014a). Differences between Presentation Methods in Working Memory Procedures: A Matter of Working Memory Consolidation. Journal of Experimental Psychology. Learning, Memory, and Cognition, 40(2), 417–428. 10.1037/a0034301

Ricker, T. J., Nieuwenstein, M. R., Bayliss, D. M., & Barrouillet, P. (2018). Working memory consolidation: Insights from studies on attention and working memory. Annals of the New York Academy of Sciences, 1424(1), 8–18. 10.1111/nyas.13633

Scheeringa, R., Petersson, K. M., Oostenveld, R., Norris, D. G., Hagoort, P., & Bastiaansen, M. C. M. (2009). Trial-by-trial coupling between EEG and BOLD identifies networks related to alpha and theta EEG power increases during working memory maintenance. NeuroImage, 44(3), 1224–1238. 10.1016/j.neuroimage.2008.08.041

Schneider, D., Barth, A., & Wascher, E. (2017). On the contribution of motor planning to the retroactive cuing benefit in working memory: Evidence by mu and beta oscillatory activity in the EEG. Neuroimage, 162, 73–85. 10.1016/j.neuroimage.2017.08.057

Souza, A. S., & Oberauer, K. (2017). Time to process information in working memory improves episodic memory. Journal of Memory and Language, 96, 155–167. 10.1016/j.jml.2017.07.002

Thaler, L., Schütz, A. C., Goodale, M. A., & Gegenfurtner, K. R. (2013). What is the best fixation target? The effect of target shape on stability of fixational eye movements. Vision Research, 76, 31–42. 10.1016/j.visres.2012.10.012

Unsworth, N. (2016). Working memory capacity and recall from long-term memory: Examining the influences of encoding strategies, study time allocation, search efficiency, and monitoring abilities. Journal of Experimental Psychology: Learning, Memory, and Cognition, 42(1), 50–61. 10.1037/xlm0000148

Van Ede, F. (2018). Mnemonic and attentional roles for states of attenuated alpha oscillations in perceptual working memory: A review. European Journal of Neuroscience, 48(7), 2509–2515. 10.1111/ejn.13759

van Wijk, B. C. M., Daffertshofer, A., Roach, N., & Praamstra, P. (2009). A Role of Beta Oscillatory Synchrony in Biasing Response Competition? Cerebral Cortex, 19(6), 1294–1302. 10.1093/cercor/bhn174

Wen, W., Grover, S., Hazel, D., Berning, P., Baumgardt, F., Viswanathan, V., … Reinhart, R. M. G. (2024). Beta-band neural variability reveals age-related dissociations in human working memory maintenance and deletion. PLOS Biology, 22(9), e3002784. 10.1371/journal.pbio.3002784

Zhuang, P., Toro, C., Grafman, J., Manganotti, P., Leocani, L., & Hallett, M. (1997). Event-related desynchronization (ERD) in the alpha frequency during development of implicit and explicit learning. Electroencephalography and Clinical Neurophysiology, 102(4), 374–381. 10.1016/S0013-4694(96)96030-7

